# Differentially regulated orthologs in sorghum and the subgenomes of maize

**DOI:** 10.1101/120303

**Authors:** Yang Zhang, Daniel W. Ngu, Daniel Carvalho, Zhikai Liang, Yumou Qiu, Rebecca L. Roston, James C. Schnable

## Abstract

Cross-species comparisons of transcriptional regulation have the potential to identify functionally constrained transcriptional regulation and genes for which a change in transcriptional regulation correlates with a change in phenotype. Conventional differential gene expression analysis and a different approach based on identifying differentially regulated orthologs (DROs) are compared using paired time course gene expression data from two species which respond similarly to cold – maize (*Zea mays*) and sorghum (*Sorghum bicolor*). Both approaches suggest that, for genes conserved at syntenic positions for millions of years, the majority of cold responsive transcriptional regulation is species specific, although initial transcriptional responses to cold appear to be more conserved between the two species than later responses. In maize, the promoters of genes with both species specific and conserved transcriptional responses to cold tend to contain more micrococcal nuclease hypersensitive sites in their promoters, a proxy for open chromatin. However, genes with conserved patterns of transcriptional regulation between the two species show lower ratios of nonsynonymous to synonymous substitutions consistent with this population of genes experiencing stronger purifying selection. We hypothesize that cold responsive transcriptional regulation is a fast evolving and largely neutral molecular phenotype for the majority of genes in Andropogoneae, while a smaller core set of genes involved in perceiving and responding to cold stress are subject to functionally constrained cold responsive regulation.

## Introduction

The grasses are a clade of more than 10,000 species, which exhibit conserved morphology and genome architecture (Bennetzin and Freeling, 1993). Grasses have adapted to grow in a wide range of climates and ecologies across the globe, with 20% of total land area covered by ecosystems dominated by grasses (Shantz, 1954). As a result, the range of tolerance to abiotic stresses present in the grass family (Poaceae) far exceeds that present within any single grass species. However, to date, studies attempting to identify determinants of abiotic stress tolerance at a genetic or genomic level have predominantly focused on individual species (Chopra *et al*., 2017; Priest *et al*., 2014; Revilla *et al*., 2016; Tiwari *et al*., 2016; Waters *et al*., 2017). Identifying changes in gene regulation across even closely related species is more challenging, and the associated methods are far less advanced. Here we seek to develop effective methods for comparing gene regulatory patterns between syntenic orthologous genes in closely related species.

The majority of genetic changes with phenotypic effects can be broadly classified into two categories, those that alter protein coding sequence, and those that alter the regulation of gene expression. DNA sequence changes that alter protein coding sequence can be identified in a straightforward fashion. The probability that a given polymorphism in protein coding sequence will have a phenotypic effect can also often be estimated. At a basic level this involves classification as synonymous, missense and nonsense mutations. However, information on the overall level of evolutionary conservation for a given amino acid residue can also be used to increase the accuracy of these predictions (Cooper *et al*., 2005; Ng and Henikoff, 2001; Reva *et al*., 2011). Both genes involved in development, and those involved in abiotic stress response tend to be associated with large quantities of conserved regulatory sequences (Freeling *et al*., 2007; Sun *et al*., 2010; Turco *et al*., 2013). Single species studies have revealed that abiotic stresses in general cause changes in regulation of large proportions (10-20%) of expressed genes (Chopra *et al*., 2015; Lee *et al*., 2005; Makarevitch *et al*., 2015; Venu *et al*., 2013).

For initial cross species comparisons, data on changes in the transcriptional responses to cold stress of maize and sorghum was employed. Cold was selected as a stress which could be delivered in a consistent fashion and time frame. Maize and sorghum were selected based on their close evolutionary relationship (Swigoňová *et al*., 2004), high quality sequenced genomes (Paterson *et al*., 2009; Schnable *et al*., 2009), and common susceptibility to cold stress (Chinnusamy *et al*., 2007; Hetherington *et al*., 1989; Wendorf *et al*., 1992). One ultimate goal of cross species comparisons of transcriptional regulation would be to link changes in regulation to changes in phenotype. Initially, comparing closely related species where gene regulation should be largely conserved provides a more useful platform for developing and testing comparative approaches. Comparison of syntenic orthologous genes in maize and sorghum indicated both conserved and lineage specific patterns of cold responsive regulation are common, with different biological functions represented in these two classes of cold-responsive genes. Conserved patterns of cold responsive regulation are more common initially, however as the length of cold stress increases, the number of genes with conserved responses in both species is not significantly greater than what would be observed in random data. Interestingly, chromatin accessibility in maize plants grown under non-stressed conditions, as measured using micrococcal nuclease hypersensitivity (MNase HS) (Rodgers-Melnick *et al*., 2016; Vera *et al*., 2014), showed distinct patterns in genes which respond to cold only in maize compared to genes which respond to cold in sorghum but not in maize.

## Results

Parallel analyses were conducted using a set of 15,232 syntenic orthologous gene pairs conserved between the maize1 subgenome and sorghum and 9,554 syntenic gene pairs conserved between the maize2 subgenome and sorghum (Schnable *et al*., 2012; Tang *et al*., 2011). Gene expression data generated from whole seedlings under control conditions showed syntenic orthologs exhibited reasonably well correlated patterns of absolute gene expression levels between sorghum and either subgenome of maize (Spearman’s rho =0.79-0.84, Pearson r = 0.67-0.85, Kendall rank correlation 0.67-0.63, supplementary fig. S1). This observation is consistent with previous reports from analysis of expression across reproductive tissues in three grass species (Davidson *et al*., 2012).

The lethal effect of prolonged cold stress on maize and sorghum (Ercoli *et al*., 2004; Hetherington *et al*., 1989; Olsen *et al*., 1993; Sánchez *et al*., 2014; Shaykewich, 1995) could be visually confirmed following prolonged cold treatment (fig. 1A–C, supplementary fig. S2, See Methods). Measurements of impairment of CO_2_ assimilation rates after recovery from a controlled length cold stress were employed to provide more quantitative measures of cold stress tolerance. Six panicoid grass species were assessed using this method (fig. 1D). After one day of cold stress, the species can be broadly classified as either cold stress insensitive or cold stress sensitive with both maize and sorghum in the cold stress sensitive category. A longer period of cold stress (3 days) demonstrated greater impairment of CO_2_ assimilation rates in sorghum than in maize, consistent with previous reports on the relative cold sensitivity of these two species (Chinnusamy *et al*., 2007; Chopra *et al*., 2017; Fiedler *et al*., 2016; Hetherington *et al*., 1989; Wendorf *et al*., 1992).

**FIG. 1.**
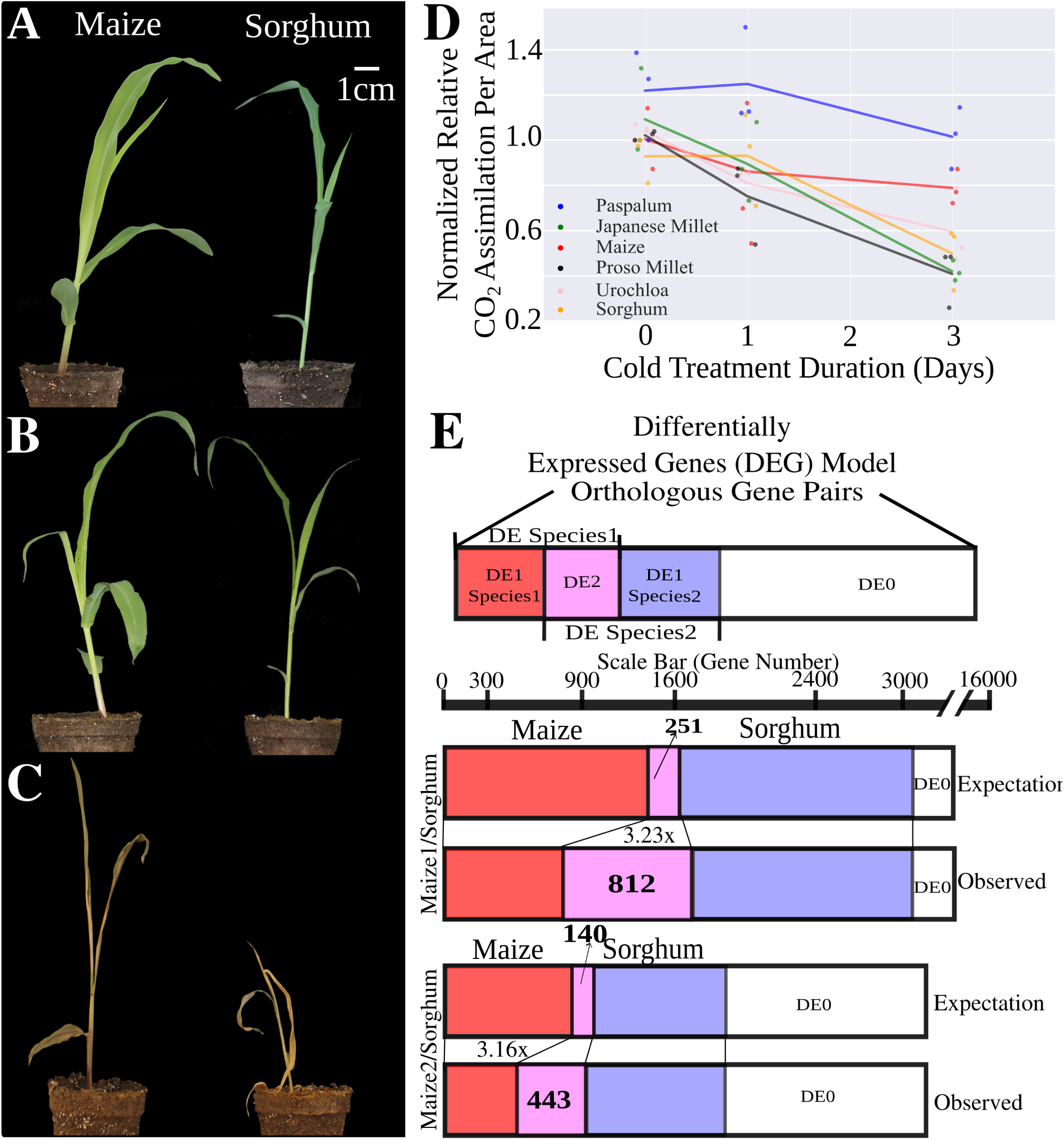
(A-C) Representative seedling phenotypes for maize (left) and sorghum (right). (A) control conditions (B) 24 hours of stress at 6 degrees Celsius (C) 14 days at 6 degrees Celsius and two days recovery under greenhouse conditions. (D) Normalized relative CO_2_ assimilation rates for plants measured under control conditions or after 1 or 3 days of cold stress. Individual datapoints were jittered on the x-axis to avoid overlap and improve readability. (E) Comparison of expected and observed value for gene pairs identified as differentially expressed in response to cold in zero, one, or both species using either maize1/sorghum or maize2/sorghum pairs. Expected distributions were calculated based on a null hypothesis of no correlation in gene regulation between maize and sorghum (see Methods).

### Conventional differentially expressed gene analysis

Cold responsive gene expression was first quantified by comparing gene expression after one day of cold stress – when maize and sorghum still exhibit comparable levels of CO_2_ assimilation impairment (fig. 1D)– to control samples collected prior to the initiation of cold stress (supplementary table S1). Among maize1/sorghum syntenic gene pairs 1,634 (10.7%) and 2,343 (15.4%) genes were classified as differentially expressed genes (DEGs) between control and cold stress in each species, respectively, using conventional differential gene expression analysis (see Methods). For maize2/sorghum syntenic gene pairs these values were 927 (9.7%) and 1,446 (15.1%) genes, respectively. Only 812 (5.3%) of maize1/sorghum syntenic genes were classified as showing a differential regulation in response to cold in both species (fig. 1E). This is greater than the 251 genes pairs expected if cold responsive gene regulation was not correlated between the two species, but indicates that a maximum of approximately three quarters (74.3%) of genes identified as responding to cold in both species do so as a result of common descent from an ancestral cold responsive gene in the common ancestor of maize and sorghum. Extending this calculation to the total group of genes identified as transcriptionally responding to cold in maize1 or sorghum only 34.1% and 34.7% respectively are calculated to have retained a conserved pattern of cold responsive gene expression since the divergence of maize and sorghum from a common ancestor 12 million years ago (Swigoňová *et al*., 2004). Results for maize2/sorghum gene pairs were comparable.

Given the close relationship of these two species (Swigoňová *et al*., 2004), the fact that both are indigenous to tropical latitudes (De Wet, 1978; van Heerwaarden *et al*., 2011), the high degree of promoter conservation observed in abiotic stress responsive genes (Freeling *et al*., 2007), the apparent low degree of conservation in cold stress responsive relation should have been unexpected. However, while being conceptually unexpected, this result is consistent with studies that have found significant divergence in abiotic stress responses between different haplotypes in maize (Waters *et al*., 2017)

One potential explanation is that the same cold stress pathways are being induced in maize and sorghum, but these pathways are induced more rapidly in one crop than the other when exposed to equivalent cold stresses. To test this potential explanation, a more detailed time course, comparing expression levels between matched pairs of cold stressed and control plants of each species at six time points distributed over 24 hours was employed (See Methods and supplementary table S1). The proportion of genes classified as differentially expressed at different time points ranged from 0.03 to 0.42 for maize1/sorghum gene pairs and 0.03 to 0.38 for maize2/sorghum gene pairs. Comparing the number of genes identified as differentially expressed in each of all 36 possible pairwise combinations of time points between the two species showed that the greatest proportion of shared differentially expressed gene pairs was identified when identical time points were compared between the two species and that the overall number of shared differentially expressed gene pairs increases at later time points (fig. 2A). However, after controlling for the number of shared differentially expressed gene pairs if gene regulation was uncorrelated between the two species, as described above, early time points show much higher expected proportions of shared differentially expressed genes resulting from common descent from an ancestral cold responsive gene (fig. 2B).

**FIG. 2.**
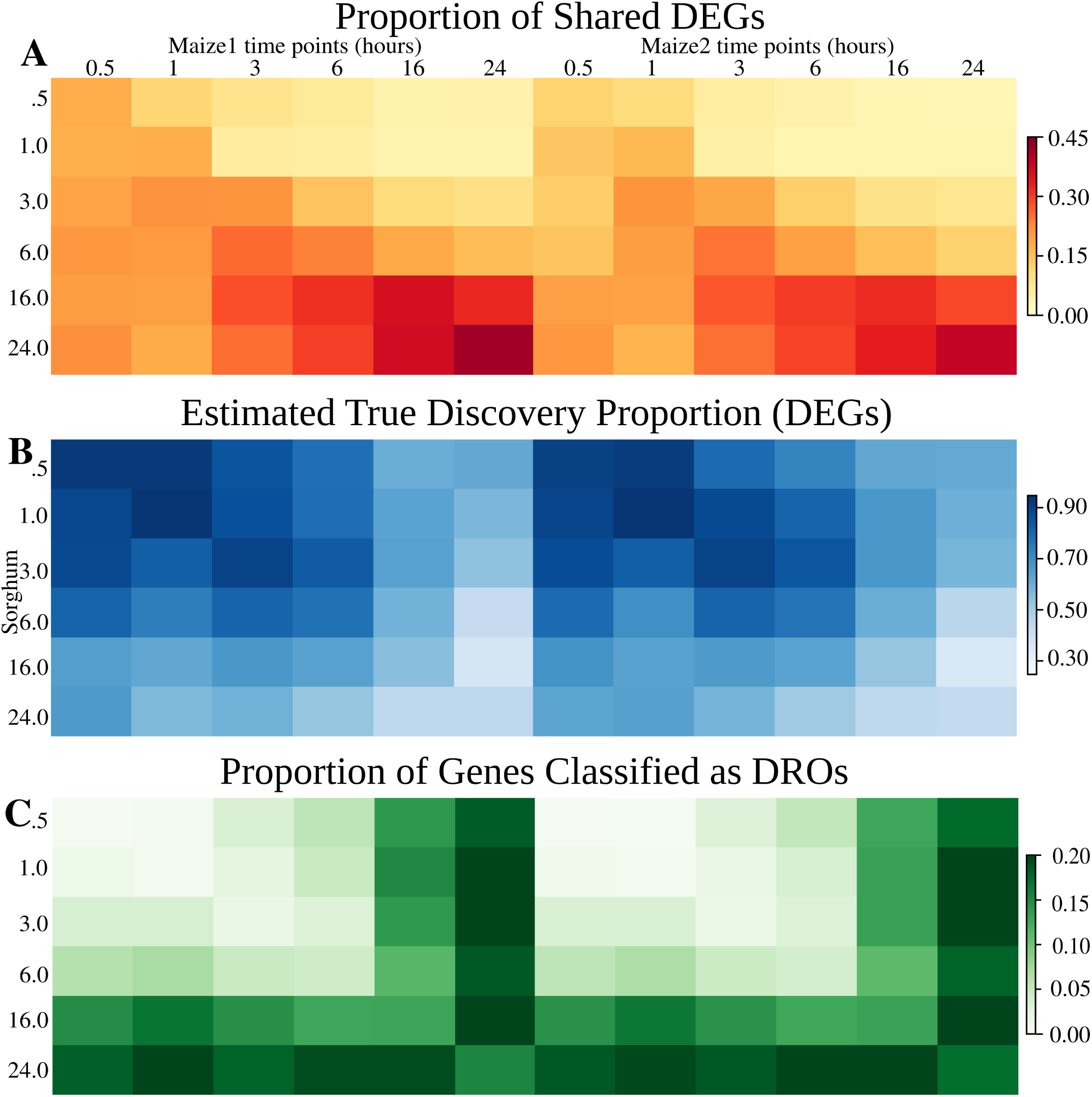
(A) Proportion of gene pairs identified as DEGs in either maize or sorghum which were identified as DEGs in both maize and sorghum (ie DE2 genes). (B) Estimated true discovery percentage for DE2 genes (observed DE2 genes - expected DE2 genes)/observed DE2 genes) using the same null model described in fig. 1A. (C) Proportion of gene pairs classified as DROs on in each of 36 possible comparisons between a sorghum time point and a maize time point.

### Differentially regulated ortholog analysis

The conceptually unexpected but that relatively few shared differentially expressed genes were identified between maize and sorghum has another potential explanation. Differential gene expression analysis may not be testing the correct null hypothesis for between dataset or between species comparisons (Paschold *et al*., 2014). The null hypothesis of conventional DEG analysis is that the expression values observed for a given gene under control and stress conditions are drawn from the same underlying distribution. This approach is perfectly suitable for single species analysis. In a two-species analyses, such as those conducted above, a DEG approach divides gene pairs into three categories: genes pairs classified as differentially expressed in neither species (DE0), in one species but not the other (DE1) and in both species (DE2, fig. 3A).

**FIG. 3.**
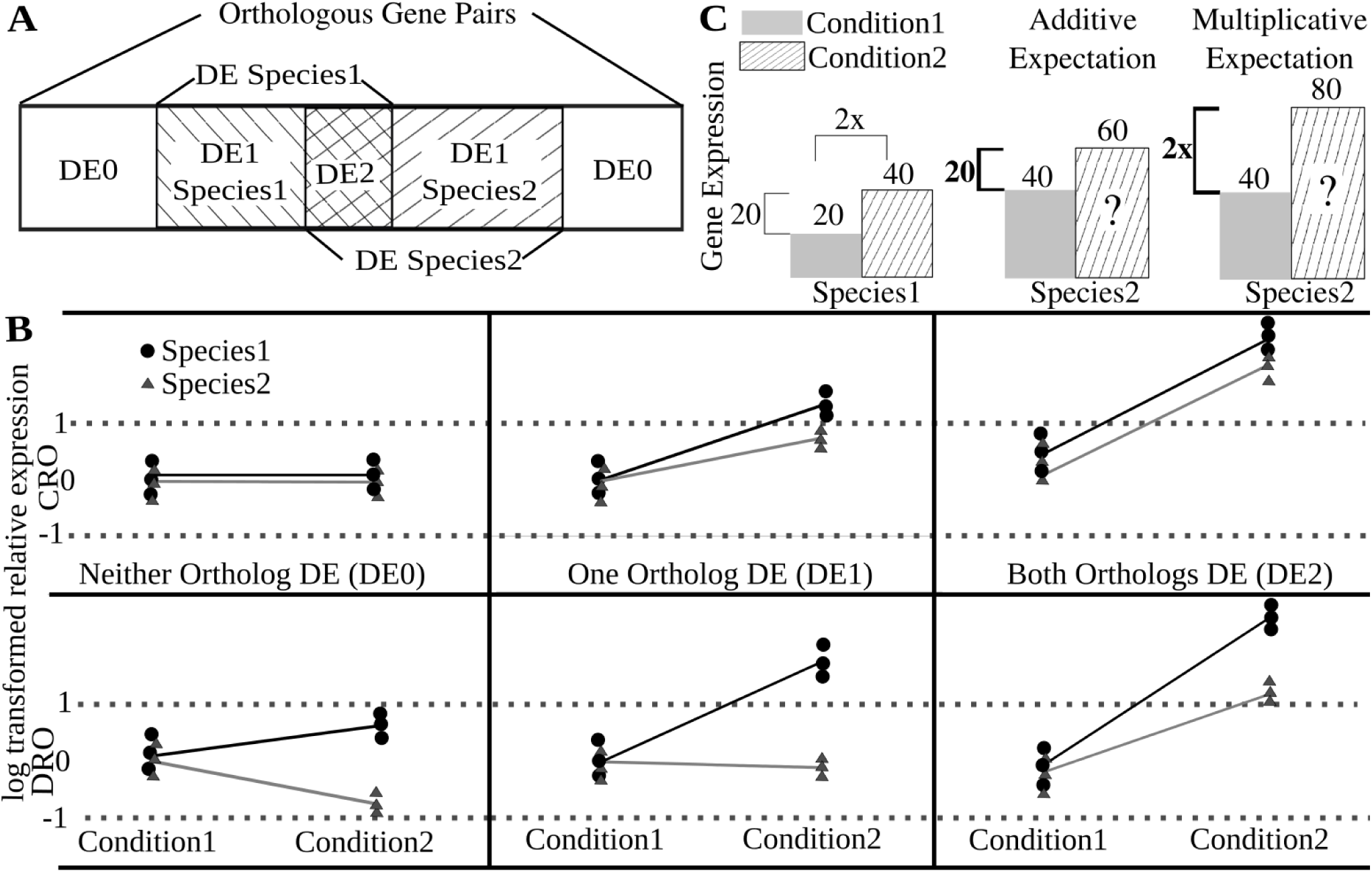
(A) Illustration of DEG based gene pair classification model. (B) Illustration of the different classification results which can be produced for a given gene pair using either a DEG-based analysis – testing whether each gene changes significantly between conditions – or a DRO-based analysis – testing whether the pattern across the two conditions is significantly different between copies of the same gene in the two species. (C) Two models – additive and multiplicative – for predicting what the same pattern of gene regulation should look like when the underlying level of expression changes.

As shown in fig. 3B, in principle, each of those three categories can include gene pairs without significant differences in the pattern of regulation between species (comparably regulated orthologs or CROs), as well as gene pairs that do show significant differences in regulation between the two species (differentially regulated orthologs or DROs). Distinguishing between DROs and CROs requires testing a different null hypothesis: that the change in expression for a given gene between two treatments is equivalent to the change in expression for an ortholog of that same gene, in a different species, across the same two treatments. Testing this null hypothesis across species in turn requires defining an accurate model of what the same pattern of gene expression looks like in starting from different baseline levels of expression.

For an orthologous gene pair which is expressed at different baseline levels in two species, there are two, different, models which can be used to compare a change in expression between treatment and control: additive and multiplicative (fig. 3C). When expression under control conditions is equivalent between the two species, these models yield the same predicted expression under stressed conditions. However, when control condition expression is different between the two species, the models produce different expected expression values under stress conditions.

To test which of these models is a better representation of how cold responsive gene regulation actually operates in maize, a set of 5,257 gene pairs retained from the maize whole genome duplication (WGD) (Schnable *et al*., 2011) was utilized. The maize whole genome duplication created two copies of each gene in the genome, each associated with the same chromatin environments and regulatory sequences. The expression level of WGD gene pairs in the maize genome in the same samples results from the same trans-factors acting in exactly the same tissue and cell types, so divergence in the regulation of these genes should result only from cis-regulatory variation which has accumulated since the maize WGD (Freeling *et al*., 2012).

To test the additive and multiplicative null models, the expression pattern of one maize gene copy between control and cold stress was used to predict the expression pattern of the other maize gene copy using each null model from fig. 3C. Analysis was conducted in parallel at each of the six time points in maize using maize1/maize2 gene pairs where at least one copy was identified as differentially expressed at that time point. Gene pairs were omitted from the analysis if the predictions of both models were more similar to each other than either was to the observed value.

The multiplicative model was more accurate at predicting cold responsive expression patterns between maize WGD duplicates than the additive model at all time points (supplementary table S2). Requiring the difference between the predictions of the two models be at least twice as large as the difference between the better model and the observed expression pattern produced similar results (supplementary fig. S3, supplementary table S2).

Genes were classified as DROs or CROs using DESeq2’s test for interaction between multiple factors (Love *et al*., 2014) (See Methods). Fig. 2C shows the proportion of gene pairs classified as DROs in each 36 possible pairwise combination of maize and sorghum time points. Consistent with the earlier DEG analysis, fewer gene pairs show significant differential regulation between the two species at early time points and more gene pairs show significant differential regulation at later time points. Comparing the same time points for maize and sorghum identifies fewer differentially regulated orthologs than comparisons between non-equivalent time points in the two species. Fewer differentially regulated orthologs were identified at earlier cold treatment time points than later long cold treatment period. This is consistent with the analysis of DEGs above which suggested early cold stress responses were more conserved across sorghum and maize than later cold stress responses. Notably, DRO analysis produces the opposite conclusion from a naive interpretation of DEG analysis of the same dataset. The latter indicated that more gene pairs are differentially expressed in both species at later time points, while the former demonstrates that the transcriptional responses of maize and sorghum to cold stress show increased divergence at later time points.

### Functional differences between genes with conserved or lineage specific regulatory patterns

Genes classified as responding to cold stress in both species (DE2) tended to have significantly lower ratios of nonsynonymous nucleotide changes to synonymous nucleotide changes (Ka/Ks ratio) than genes which responded to cold stress in only one species or in neither species. This indicates genes with conserved patterns of cold responsive regulation experience stronger purifying selection than genes with lineage specific patterns of cold responsive regulation (fig. 4A–B). Genes in the DE2 category were also more likely to be classified as transcription factors by Grassius (Yilmaz *et al*., 2009).

**FIG. 4.**
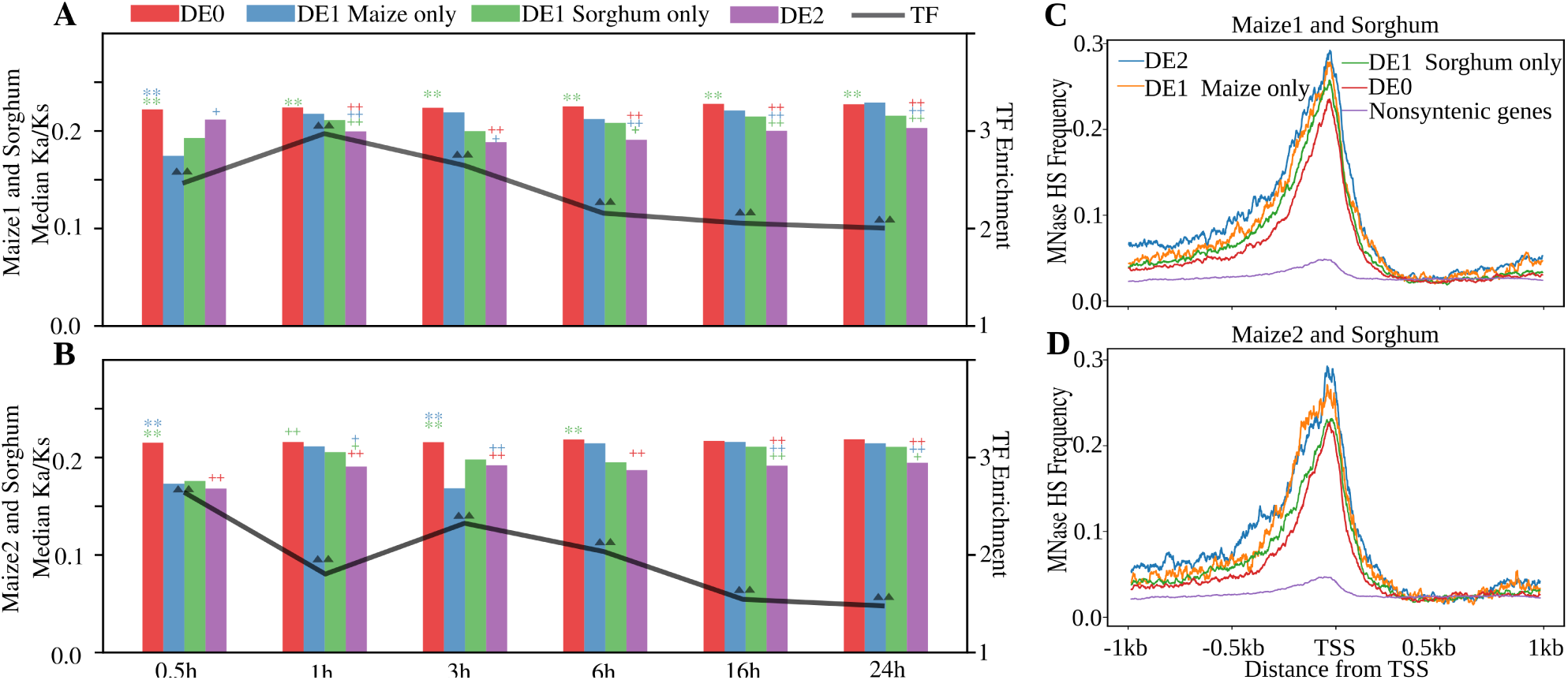
Median synonymous substitution rates between maize and sorghum for gene pairs classified as DE0, DE1, or DE2 at each of the six time points sampled. The enrichment of genes annotated as transcription factors among DE2 gene pairs at the same time points relative to all gene pairs is shown with the overlaying black line. Statistical comparisons between DE0 vs DE1 Maize only, DE0 vs DE1 sorghum only, and DE0 vs DE2 categories performed using a t-test. “*” with the same color as bars. *: p-value<0.05; **: p-value<0.01. Statistical comparisons between DE2 vs DE1 Maize only and DE2 vs DE1 sorghum only categories, “+” with the same color as bars. +: p-value<0.05; ++: p-value<0.01; Statistical comparisons between DE0 vs all the orthologous syntenic gene pairs category performed using Fisher’s exact test, Δ: p-value<0.05; ΔΔ: p-value<0.01. (A) Results for maize1/sorghum gene pairs. (B) Results for maize2/sorghum gene pairs. Association between gene pair expression pattern at the 24 hour cold stress time point and open chromatin – using a MNase hypersensitivity assay (Rodgers-Melnick *et al*., 2016) – measured in maize seedlings under control conditions. (C) Results for maize1/sorghum gene pairs (D) Results for maize2/sorghum gene pairs.

Finally, chromatin states in the promoters of genes with different patterns of cold responsive regulation were examined using a published dataset of MNase hypersensitive sites generated from maize seedlings grown under non-stressed conditions (Rodgers-Melnick *et al*., 2016). Comparisons were made for maize DE0, Maize DE1, Sorghum DE1, DE2 and nonsyntenic genes at each of the six cold stress time points. Many nonsyntenic genes responded to cold, however nonsyntenic genes as a whole showed little or no open chromatin as defined by MNase HS associated with their TSS (transcription start site) or proximal promoters. Previous studies of other epigenetic marks have also concluded that the chromatin signals of nonsyntenic genes in maize are more similar to intergenic sequence than to syntenic genes (Eichten *et al*., 2011). All categories of syntenic genes tended to have a peak of MNase sensitivity associated with their TSS and more open chromatin in their proximal promoters than nonsyntenic genes. Genes with conserved cold responsive regulation (DE2) appear have the greatest amount of open chromatin in their proximal promoters (fig. 4C–D). Intriguingly, the maize copies of maize DE1 gene pairs exhibited stronger open chromatin signals that the maize copies of sorghum DE1 gene pairs even though data on MNase hypersensitive sites came from seedlings grown under control conditions. Similar patterns were observed using analysis of each of the six cold stress time points, and were robust to the use of minimum expression cut offs (supplementary fig. S4).

## Discussion

Previous studies in maize and sorghum have reported that 10-15% of all expressed genes respond transcriptionally to cold stress (Chopra *et al*., 2015; Makarevitch *et al*., 2015). A study of three genotypes of maize using F1 hybrids reported that between 1.7-14.6% of genes assayed and 4.2-33.2% of cold responsive genes showed significant cis-regulatory variation between alleles in response to cold stress (Waters *et al*., 2017). Here we found that, while a consistent percentage of conserved genes responded transcriptionally to cold stress in maize and sorghum, the proportion of genes which responded similarly between these two species was much lower. Correcting for the expected overlap across conserved genes based solely on the proportion of genes exhibiting cold responsive transcriptional changes in each species further reduced the expected number of gene pairs where shared regulation resulted from the conservation of an ancestral pattern of cold responsive transcriptional regulation. These data imply that cold stress responsive regulation is a relatively fast evolving trait in panicoid grasses, a result that has significant implications for the interpretation of stress response RNA-seq datasets. We demonstrated that genes which are respond to cold only in a single lineage experience lower levels of purifying selection and are less likely to be annotated as transcription factors.

It appears that a relatively small core set of genes exhibit conserved responses to cold across the two species in this initial analysis. Thus, we propose a model where a small core set of genes involved in the mechanisms by which panicoid grasses perceive and respond to cold stress are under functionally constrained cold responsive transcriptional regulation, while a much larger set of genes can gain or lose cold responsive transcriptional regulation in a neutral fashion, potentially through transposon mediated mechanisms (Makarevitch *et al*., 2015; Naito *et al*., 2009). Consistent with this model, the genes with conserved cold responsive gene regulation were observed to have lower ratios of nonsynonymous to synonymous coding sequence substitions, which would imply their coding sequence is also subject to greater functional constraint. This model would also be consisent with the relatively high proportion of maize cold responsive genes which exhibit variation in cold responsive regulation across alleles (Waters *et al*., 2017).

### The challenge of linking genes to functions based on expression evidence

The model above would predict than the observation of stress-responsive changes in transcript abundance in a single species are not strong evidence that the associated gene plays a role in the response to that particular stress. While sequencing genomes and identifying genes is becoming a more straightforward task, confidently assigning functional roles to newly identified genes remains challenging. Of the 34,211 genes in sorghum, 5,526 genes (16.2%) are not associated with any GO terms in the latest release of phytozome. The numbers are even lower in maize, which has 22,249 out of 63,480 genes (35.1%) without any GO terms in the latest release of phytozome. Current functional annotation methods are based primarily on homology, which are more effective at defining the molecular functions of proteins than their role in whole organism processes.

While transcriptional response in a single species may not be a strong link to a functional role, it may prove to be the case that functionally constrained transcriptional responses are an effective method for identifying these links. Collecting parallel expression datasets in multiple species can be time consuming and costly. A number of alternative approaches to identifying functionally constrained cold responsive transcriptional regulation were tested above. Early transcriptional responses to cold (30 minutes-3 hours) appeared to show greater conservation across species than later transcriptional responses. Regions of open chromatin detected through MNase hypersensitivtiy (Rodgers-Melnick *et al*., 2016; Vera *et al*., 2014), were preferentially associated with genes which responded transcriptionally to cold stress in maize, however, this association was observed for genes with either conserved or lineage specific patterns of cold responsive regulation.

### Importance of developing methods for cross-species comparisons of transcriptional regulation

Both modeling (Orr, 1998, 1999) and empirical studies (Chan *et al*., 2010; Studer *et al*., 2011) have found that genetic variants responsible for large sudden changes in natural or artificial selection tend to have large and pleiotropic effects. In maize, distinct genetic architecture underlying traits which had been subjected to selection during domestication – one large effect QTL and many small modifiers – and traits which were not selected on during domestication – many small effect QTL (Wallace *et al*., 2014). This model was supported by recent work with an inter-subspecies cross of maize and its wild progenitor teosinte (*Zea mays* ssp. *parviglumis*). Looking at tassel morphology, distinctly genetic architectures were reported for traits believed to have been under selection during domestication compared to those traits which were not (Xu *et al*., 2017). Developing effective approaches for comparing transcriptional regulation of conserved syntenic genes across related grass species has the potential to identify large effect polymorphisms responsible for interspecies phenotypic variation in traits such as abiotic stress tolerance where substantial phenotypic variation exists between species (fig. 1D).

Here we have shown that by using synteny to identify pairs of conserved orthologs across related species, it is possible to identify species by treatment interactions signifying changes in gene regulation across species (DROs) using a multiplicative model of gene regulation. The use of a multiplicative model was in turn supported by analysis of the regulation of duplicated maize genes within the same sample. By increasing the number of species sampled, it may soon be possible to define a consistent core set of genes subjected to functionally constrained regulation in response to cold across the grasses. Changes in the regulation of these core genes in specific lineages with different cold stress response phenotypes would be useful candidates for the type of large effect changes predicted to produce between species phenotypic variation. However, the interpretation of such data must take into account that, unlike within species studies of allelic variation in in cold responsive regulation, between species analysis can not distinguish cis-regulatory and trans-regulatory sources of variation in transcriptional responses.

## Materials and Methods

### Plant growth and cold treatment

For maize and sorghum we employed the reference genotypes used for genome sequencing and assembly B73, and BTx623, respectively. SNP calling using RNA-seq data from B73 was used to verify that the plants used in this study came from the USA South clade of B73 accessions, those closest to the original reference genome (Liang and Schnable, 2016). Under the growing conditions employed, maize developed faster than sorghum, and sorghum seedlings twelve days after planting (DAP) were selected as being roughly developmentally equivalent to 10 DAP maize seedlings based on leaf number and morphology (fig. 1A). Planting dates were staggered so that all species reached this developmental time point simultaneously. For the original RNA-seq presented in fig. 1A, seeds were planted in metroMix 200 and grown in greenhouse conditions under 13 hours day time in greenhouses of University of Nebraska-Lincoln’s Beadle Center, with target conditions of 320 mol m^−2^ s^−1^, 13 hours/11 hours 29°C/23°C day/night and 60% relative humidity. Control plants were harvested directly from the greenhouse three hours before lights off. Plants subjected to cold stress treatment were moved to a cold treatment, with 33 mol m^−2^ s^−1^, 12 hours/12 hours 6°C/6°C day/night. Cold stressed plants were harvested three hours before lights off. Each sample consisted of pooled above ground tissue from at least three seedlings. Each biological replicate was harvested from plants that were planted, grown, and harvested at a distinct and separate time from each other biological replicate. For the time course RNA-seq presented in Figure 2 and onward in the study, maize and sorghum were planted as above, and grown in a Percival growth chamber with target conditions of 111 mol m^−2^ s^−1^, 60% relative humidity, a 12 hour/12 hour day night cycle with a target temperature of 29° C during the day and 23° C at night. Onset of cold stress was immediately before the end of daylight illumination at which point half of the plants were moved to a second growth chamber with equivalent settings with the exception of a target temperature of 6°C both during the day and at night. Each sample represents a pool of all above ground tissue from at least three seedlings. Samples were harvested from both the paired control and cold stress treatments at 0.5 hours, 1 hour, 3 hours, 6 hours, 16 hours, and 24 hours after the onset of cold stress. Biological replicates included both maize and sorghum plants where were offset in planting but stressed and harvested at the same time in the same growth chambers.

### CO_2_ assimilation rates measurement

Plants were grown and cold treated as above, with the modification that in the cases of sorghum, small plastic caps were placed over the seedlings to prevent plants from becoming too tall to fit into the LiCor measurement chamber (approximately two inches). After 0, 1, or 3 days of cold treatment, plants were allowed to recover in the greenhouse overnight. The following morning, CO_2_ assimilation rates were measured using the the Li-6400 portable photosystem unit under the following conditions: PAR 200 mol mol^−1^, CO_2_ at 400mol mol^−1^ with flow at 400 mol mol^−1^ and humidity at greenhouse conditions. Whole plant readings were measured for sorghum, foxtail millet, paspalum, and urochloa after covering their pots with clay and using the LiCor “Arabidopsis chamber.” Maize was measured using the leaf clamp attachment which was consistently placed on the second leaf at a position 3 cm above the ligule. Leaf area was measured using the Li-3100c Area meter (Li-Cor). Accessions used for each species presented in fig. 1D were: paspalum (*Paspalum vaginatum*): 540-79, Japanese millet (*Echincloa esculenta*): USDAPI 647850, proso millet (*Panicum miliaceum*): earlybird USDAPI 578073, urochloa (*Urochloa fusca*): common name AB4, sorghum: BTx623, and maize: B73.

### Identifying syntenic orthologs

Coding sequence data for primary transcripts of each annotated gene in the genome assemblies of 8 grass species including maize and sorghum in the analysis was obtained from Phytozome 10.2. Similar sequences were identified using LASTZ (Harris, 2007), requiring an alignment spanning at least 50% of total sequence length and 70% sequence identity. In addition, the arguments –ambiguous=iupac, –notransition, and–seed=match12 were all set in each run. LASTZ output was converted to QuotaAlign’s “RAW” format using a version of the blast_to_raw.py script which had been modified to take into account differences in output format between BLAST and LASTZ. The additional parameters –tandem_Nmax=10 and –cscore=0.5 were specified when running this script.

RAW formatted data was processed using the core QuotaAlign algorithm with the parameters–merge, and –Dm=20. –quota was set to 1:2 in comparisons to maize and 1:1 in all other comparisons. Pure QuotaAlign pan-grass syntenic gene sets were constructed using this dataset directly. Polished QuotaAlign pan-grass syntenic gene sets were constructed by first predicting the expected location for a given query gene in the target genome, and then selecting the gene showing the greatest sequence similarity (as determined by lastz alignment score) within the window from 20 genes downstream of the predicted location to 20 genes upstream of the predicted location. The final syntenic gene list used here is available from the following link: figshare:10.6084/m9.figshare.3113488.v1.

### RNA-seq data generation

RNA isolation and library construction followed the protocol described by Zhang et al. (Zhang *et al*., 2015). The number of reads generated per library are summarized in (supplementary table S1). Sequencing was conducted at Illumina Sequencing Genomics Resources Core Facility at Weill Cornell Medical College. Raw sequencing data are available through the NCBI (http://www.ncbi.nlm.nih.gov/bioproject) under accession number PRJNA343268 and PRJNA344653. Adapters were removed from raw sequence reads using cutadapt version 1.6 (Martin, 2011). RNA-seq reads were mapped to genome assemblies downloaded from phytozome: v6a (*Zea mays*), v3.1 (*Sorghum bicolor*). RNA-seq reads from each species were aligned using GSNAP version 2014-12-29 (Wu and Nacu, 2010; Wu and Watanabe, 2005) and per-gene read counts were obtained using HTSeq v. 0.6.1 (Anders *et al*., 2014).

### Identifying differentially expressed genes (DEG)

Differentially expressed genes (DEGs) were identified using count data generated as described above and DESeq2 (version 1.14.0) (Love *et al*., 2014) based on a comparison of the treatment and control with adjP-value<=0.05, meaning absolute log2 of fold Change of between treat and control>=1. All expressed syntenic orthologous genes were classified into one of three categories. Three categories include genes which were classified as respond transcriptionally to cold in at least one species (DE1) (Figure1C). The remaining category includes all expressed syntenic orthologous genes which were not classified as cold responsive in any of the two species (DE0). The number of shared genes identified as differentially expressed in two species (DE2) was tested relative to the expected overlap if there was no correlation in gene regulation across species. For the time course RNA-seq, analysis was conducted as above for all 36 possible pairwise comparisons of the 6 sorghum time points and 6 maize time points.

Estimating the true discovery percentage in analyses of DE2 genes (see fig. 1E, fig. 2B), it was necessary to calculate the number of DE2 genes expected under a null hypothesis of no conservation of gene regulation. This expected number of DE2 genes was calculated using the formula (percent of gene pairs DE in species 1)*(percent of gene pairs DE in species 2)*(total number of gene pairs analyzed was used). Total number of gene pairs was fixed at 15,232 syntenic orthologous gene pairs for maize1/sorghum comparisons and 9,554 for maize2-sorghum comparisons.

### Evaluating additive model and multiplicative models of gene regulation

From a 5,257 duplicate genes retained from the maize whole genome duplication was (Schnable *et al*., 2011) (supplementary fig. S1) in each of the six time points in maize gene pairs where both copies were classified as differentially expressed in response to cold were to test both models. The expression pattern of the maize1 gene in control and cold stress, plus the expression of the maize2 gene under control conditions was used to predict the expression of the maize2 gene under cold stress using both the additive and multiplicative models defined in fig. 3. In the relaxed case, gene pairs where the two models produced predictions which were closer to each other than either was to the observed expression value of the maize2 gene under cold stress were excluded. In the stringent case, gene pairs where the two models produced predictions that were less than twice as large as the difference between the better model and the observed value were excluded. Analyses were also conducted reciprocally using data from control and cold stress in maize2 plus data from maize1 in control conditions to predict the expression of the maize1 gene under cold stressed conditions.

### Identifying differentially regulated orthologs (DRO)

Differentially regulated orthologs were identified using count data generated as described above and an interaction term for species (maize or sorghum) and treatment (cold or control).

Interaction in DESeq2 (version 1.14.0) (Love *et al*., 2014) was used for Identifying DROs analysis. Per-gene read counts were obtained as described above and used for DESeq2 interaction analysis. Species(maize and sorghum) and condition (cold and control) were considered as two factors for design in this analysis. Simulated data for comparably regulated orthologs (CROs) generated using additive and multiplicative models was used to confirm that this approach did not classify simulated CROs based on the multiplicative model as having significant species by treatment interactions. The formula used was: design ~condition+genetype+condition:genetype. Maize sorghum gene pairs with a adjP-value<=0.001 were classified as DROs, those with adjP-value>=0.05 were classified as CROs and those with intermediate p-values were disregarded.

### Calculating Ka/Ks values

“Primary Transcript only” CDS sequences for maize (v6a), sorghum (v3.1), and setaria (v2.2) were retrieved from phytozome version 12.0. CDS sequences were translated to proteins and aligned using Kalign version 2.04 (Lassmann and Sonnhammer, 2005). The protein alignment was used as a guide to create a codon level alignment of CDS sequences. The codon alignment was supplied to PAML (version 4.09) (Yang, 2007). Synonymous and nonsynonymous substitution rates were calculated independently for each branch of the tree. When both a maize1 and maize2 gene copy were present for the same syntenic gene group, alignment and substition rate calculations were conducted separately for the maize1 gene and its syntenic orthologs in sorghum and setaria and for the maize2 and the same syntenic orthologous genes. To eliminate genes with extreme Ka/Ks ratios resulting from very low numbers of synonymous substitutions, only Ka/Ks ratios from genes with an estimated synonymous substitution rate greater than or equal to 0.05 – approximately 1/2 the median Ks ratio observed between maize and the most common recent ancestor of maize and sorghum – were considered.

### MNase hypersensitive site analysis

Intervals defined as MNase hypersenstive sites (MNase HS) were taken from (Rodgers-Melnick *et al*., 2016). Average coverage of MNase HS was calculated on a per-base basis from 1kb upstream of the annotated transcription start site (TSS) to 1 kb downstream of the transcription start site. When multiple transcripts with different TSS were present, the transcript with the earliest TSS was selected for analysis.

### Transcription Factor Enrichment Calculations

Transcription Factors (TF) enrichment was calculated using the maize transcription factor list from Grassius (Yilmaz *et al*., 2009)

## Supplementary Material

supplementary fig. S1. (A) Venn diagram showing the overlap between syntenic orthologous gene pairs conserved between maize1/sorghum and maize2/sorghum based on sharing the same sorghum co-ortholog. (B) Correlation of average expression level (log2 transformed FPKM) between maize1/sorghum and maize2/sorghum gene pairs under control conditions. (Random subsample of 1/3 of all gene pairs displayed to improve readability.)

supplementary fig. S2. Phenotypes of a larger set of seedlings under control conditions, after one day of cold treatment and after 14 days of cold treatment followed by two days recovery under greenhouse conditions.

supplementary fig. S3: Scatter plot showing the relationship between prediction error for expression under cold stress using a multiplicative model to predict expression between maize1/maize2 gene pairs or an additive model to predict expression between maize1/maize2 gene pairs. Dots in blue mark cases where the additive model was the better predictor, dots in red mark cases where the multiplicative model was the better predictor.

supplementary fig. S4: Relationship between gene pair expression pattern in maize and sorghum at each of the six time points and chromatin accessibility estimated using data on MNase HS from maize seedlings grown under control conditions. Results calculated separately for all gene pairs, and excluding gene pairs with low expression (average FPKM <2).

supplementary table S1. Number of total and aligned sequence reads per sample for both the control/stress RNA-seq experiment, and the time course RNA-seq experiment.

supplementary table S2. Number of gene pairs where the additive or multiplicative model is better predictor by absolute distance between predicted and observed values (using either standard and stringent criteria).

## Acknowledgments

This work was supported by USDA-NIFA Grant 2016-67013-24613 to RLR and JCS, NSF Award OIA-1557417 to JCS and by start-up funding from the University of Nebraska-Lincoln to RLR, YQ, and JCS.

